# Clinorotation inhibits myotube formation by fluid motion, not by simulated microgravity

**DOI:** 10.1101/2023.02.10.527979

**Authors:** Janet Mansour, Carolin Berwanger, Marcel Jung, Ludwig Eichinger, Ben Fabry, Christoph S. Clemen

## Abstract

To study processes related to weightlessness in ground-based cell biological research, a theoretically assumed microgravity environment is typically simulated using a clinostat – a small laboratory device that rotates cell culture vessels with the aim of averaging out the vector of gravitational forces. Here, we report that the rotational movement during fast clinorotation induces complex fluid motions in the cell culture vessel, which can trigger unintended cellular responses. Specifically, we demonstrate that suppression of myotube formation by 2D-clinorotation at 60 rpm is not an effect of the assumed microgravity but instead is a consequence of fluid motion. Therefore, cell biological results from fast clinorotation cannot be attributed to microgravity unless alternative explanations have been rigorously tested and ruled out. We consider two control experiments mandatory, i) a static, non-rotating control, and ii) a control for fluid motion. These control experiments are also highly recommended for other rotation speed settings and experimental conditions. Finally, we discuss strategies to minimize fluid motion in clinorotation experiments.

## Introduction

‘Simulated microgravity’ (Herranz et al., 2013) is the technical term for a ground-based laboratory condition that is used as a workaround for conducting experiments that would otherwise have to be carried out under real microgravity conditions in space. For gravity-related cell biological research, 2D-clinostats, rotating wall vessels, 3D-clinostats, random positioning machines, and magnetic levitation devices have been developed (Brungs et al., 2016; Calvaruso et al., 2021; Hasenstein, 2022; Herranz et al., 2013; Klaus, 2001; Zhang et al., 2022). The 2D-clinostat is often used because of its simplicity: cells are cultured on a flat, horizontal surface that is rotated around an in-plane axis. Cells grown along or near the horizontal axis of rotation experience negligible centrifugal forces, but are subject to a constantly changing direction of the gravitational force vector that averages out to zero. Although the rotating cells still experience the same magnitude of gravitational forces, this approach is widely considered to be a microgravity analog to conditions in space/earth orbit where the gravitational forces are balanced by centrifugal forces (Briegleb, 1992; Brungs et al., 2016; Hauslage et al., 2017; Herranz et al., 2013; Zhang et al., 2022).

Because adaptation to external forces can be rapid, especially in unicellular and small multicellular organisms, it was reasoned that the rotation of a clinostat must be sufficiently fast (Briegleb, 1992). A 2D-clinorotation speed of 60 rpm was suggested to be the most effective setting for a functional microgravity simulation (functional weightlessness) for submerged, small objects (Briegleb, 1992). Accordingly, fast clinorotation, typically with a rotational speed of 60 rpm, was then used for numerous experiments with small particles in suspension, a variety of cell types in suspension, and a few studies with adherent mammalian cells in liquid culture (Boonstra, 1999; Briegleb, 1992; Brungs et al., 2016; Cogoli, 1992; Herranz et al., 2013; Kiss et al., 2019; Moore and Cogoli, 1996). Adherent mammalian cells, for example, can be grown in plastic cell culture vessels, so-called ‘slide flasks’, that are completely filled with equilibrated medium, carefully sealed, and attached to a clinostat that rotates the vessels around the horizontal axis (Eiermann et al., 2013).

In the present work, murine C2C12 myoblasts were seeded into such slide flasks. After one day, the growth medium was replaced by a differentiation medium, and the flasks were attached to a 60 rpm-rotating 2D-slide flask-clinostat. Myotube formation was monitored over a time period of typically seven days. We found that clinorotation suppressed myotube formation, in agreement with previous work reporting reduced numbers of myotubes near the centre of rotation as well as myofibrillar changes with aberrant filamin-C and α-actinin staining patterns within these cells (Mansour Jamaleddine, 2021). The present study, however, demonstrates that this effect cannot be attributed to simulated microgravity. Instead, we show that the cellular responses are dominated by the rotation-induced fluid motion of the cell culture medium. This finding suggests that the previously reported responses of mammalian cells in liquid culture using clinostats may have been similarly dominated by rotation-induced fluid motion. We discuss strategies for improving the design of fast-rotating clinostats to minimise such fluid motion in future cell biological studies.

## Results

### Suppression of myotube formation during clinorotation does not depend on simulated microgravity

To study the effects of mechanical unloading on muscle cells, we performed 2D-clinorotation assays with differentiating C2C12 myoblasts over long periods of time (Fig. 1A-C). Visual inspection of the slide flasks by phase contrast microscopy after six to eight days of differentiation consistently showed suppressed myotube formation, which was most pronounced within an approximately 4 mm wide band with no sharp boundary along the rotational axis throughout the slide (Fig. 1D). Myotubes grown as a control on static, non-rotating slides did not show a reduced density along the centre line of the slide, but a nearly uniform distribution of myotubes (Fig. 1F). This result was initially interpreted to be related to the loss of skeletal muscle tissue observed under microgravity conditions in space (Rittweger et al., 2018), presumably because a sufficient microgravity environment was generated near the rotational axis, whereas microgravity conditions outside this narrow band break down due to above-threshold centrifugal forces (Eiermann et al., 2013).

**Figure 1.**
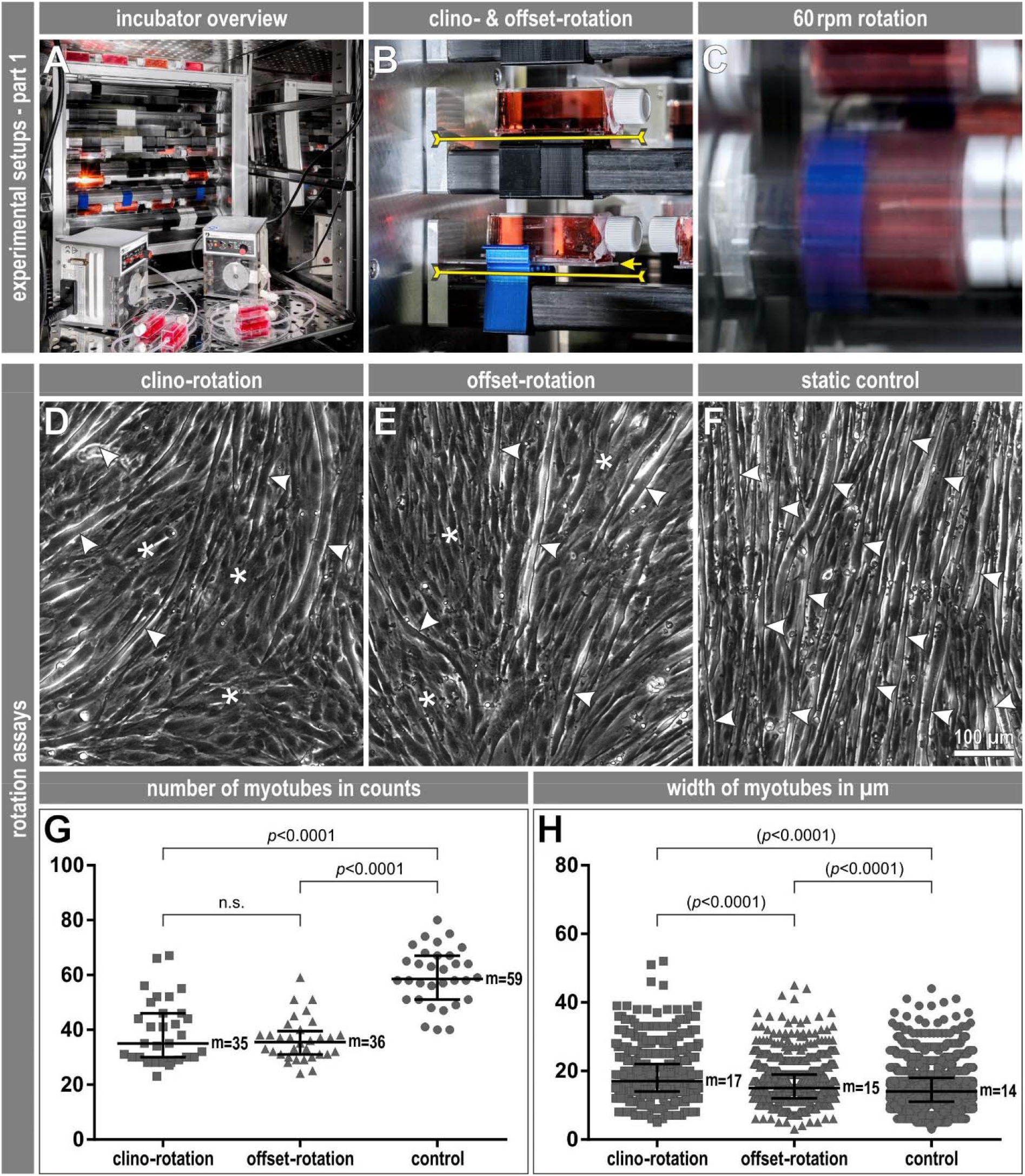
Suppression of myotube formation during clinorotation does not depend on simulated microgravity. **A**) View into the opened cell culture incubator with part 1 of the experimental setup. In the background, the slide flask 2D-clinostat with several slide flasks containing confluent layers of C2C12 myoblasts attached to the rotation axes and filled with differentiation medium. On top of the device the static, non-rotating controls. In the foreground, two peristaltic pumps with connected slide flasks converted into flow flasks by means of grommets attached to the end faces. (**B**) Detailed view of two slide flasks attached with clamps to the rotation axes of the clinostat to illustrate their usual position for clino-rotation (upper flask, black clip) compared to the additional position of offset-rotation (lower, ‘raised’ flask, blue clip) used to demonstrate side effects of the rotation in this study. Upper yellow line, position of the rotation axis in plane with the slide. Lower yellow line, offset-rotation with the rotation axis 4 mm below the slide with the adherent cells (arrow). (**C**) Slightly enlarged view of (B) during the 60 rpm fast rotation of the slide flasks. (**D-F**) Phase contrast microscopic views of C2C12 cells from the central 4 mm wide area along the rotation axis of slide flasks after 6 days of differentiation into multinucleated myotubes during (D) continuous clino-rotation, (E) continuous 4 mm offset-rotation or from (F) static, non-rotating controls. Arrowheads, examples of the formed myotubes. Asterisks, areas where myoblasts remained prominent. (**G**) Quantitation of the number of formed myotubes after 6 days of differentiation under clino-rotation, offset-rotation or static control condition. Shown is the representative result of one of a total of three independent experiments. Myoblasts of eight microscopic views from the 4 mm wide area along the central axis of rotation across the entire slide of each slide flask were counted from four slide flasks per condition of the experiment shown. Note that there is no statistically significant difference between clino-rotation and offset-rotation, but both conditions are statistically significantly different from the static, non-rotating control. (**H**) Quantification of the width of the myotubes counted in (G), each measured at the point of the largest diameter. The marginal differences in diameter are not considered biologically relevant, and therefore statistical significances are given in brackets. (G, H) Scatter dot plots with medians (values are indicated) and interquartile ranges. Statistical significances calculated by Student’s t-test subsequent to normality and homoscedasticity as well as one-way ANOVA testing.

To test the validity of the latter assumption, we mounted the slides 4 mm above the rotational axis, exposing all the adherent cells to above-threshold centrifugal forces of 0.016 RCF (Fig. 1B). Notably, this offset-rotation also led to significantly suppressed myotube formation, again within an approximately 4 mm band with no sharp boundary along the rotational axis (Fig. 1E), which was not present in the static controls (Fig. 1F) that were placed on top of the clinostat frame to expose the cells to the same vibrational forces (Fig. 1A). The number of myotubes per field of view within the 4 mm bands showed no statistically significant difference between clino-rotation (median count 35) and offset-rotation (median count 36), whereas the number of myotubes from these two conditions was statistically significantly lower compared to the static, non-rotating control (median count 59) (Fig. 1G). Myotube widths also showed statistically significant differences between groups (Fig. 1H), but these differences were small (median widths 17, 15, and 14 μm) and were not considered biologically relevant. Together, the results of the clino- and offset-rotation experiments strongly suggest that the higher myotube density outside the central band near the rotational axis is not caused by a breakdown of microgravity conditions, but must be due to previously unexplored effects. As we will show below, a plausible explanation for this observation is the presence of a rotation-induced, large cylindrical fluid motion extending over the entire length and diameter of the slide flask. The 4 mm band of suppressed myotube formation occurs where this cylinder is closest to the walls of the flask, i.e. also near the axis of rotation where fluid motion and shear stresses are largest.

### Fluid motion markedly influences C2C12 myotube formation

Previous studies reported rotation-induced motion of the cell culture medium in flasks mounted to a 3D-clinostat (two-frame rotation) (Wüest et al., 2015; Wüest et al., 2017; Zhang et al., 2022). Based on literature review and fluid dynamic simulations, a moderate rotational speed in the range of 10 rpm was suggested as being safe to avoid fluid motion-induced biological effects, although a systematic experimental study of different rotational speeds, accelerations, and induced fluid motion was lacking (Wüest et al., 2015). More extensive *in silico* analyses suggested that fluid motion can be significant even at moderate speeds, and that additional cell culture control experiments are needed to rule out fluid motion-induced effects. In particular, it was pointed out that the fluid motion within a 3D-clinorotated culture vessel does not reach a steady state, and depending on the rotational speed (6.7 rpm, 10 rpm, 15 rpm) and position within the culture vessel (wall surface versus bulk), shear stresses in a range of 10 mPa to 300 mPa can occur (Wüest et al., 2017). In contrast to 3D-clinostat rotation, theoretical studies predicted that no fluid motion occurs during steady 2D-clinorotation except during a brief acceleration phase (Klaus et al., 1998; Wüest et al., 2017).

In practice, however, the electric motor and gearbox that drive the 2D-clinostat may not deliver a perfectly constant speed, and small vibrations and rocking movements of the clinostat frame may be felt due to the off-centre mounting of the slide flasks. Therefore, we reasoned that fluid motion could also be present in 2D-clinostat experiments, and that this might influence the differentiation of myoblasts into myotubes e.g., due to small shear stresses or fluid convection that disturbs the local concentration profile of secreted auto- or paracrine differentiation signals. To test this hypothesis, we added polystyrene beads of 140, 250 or 500 μm diameter as tracer particles to medium-filled 60 rpm 2D-clinorotating slide flasks (Fig. 2A). We than mounted a small microscope objective and CCD camera to the end face of a slide flask filled with culture medium and tracer particles, and let them rotate together (Fig. 3A).

**Figure 2.**
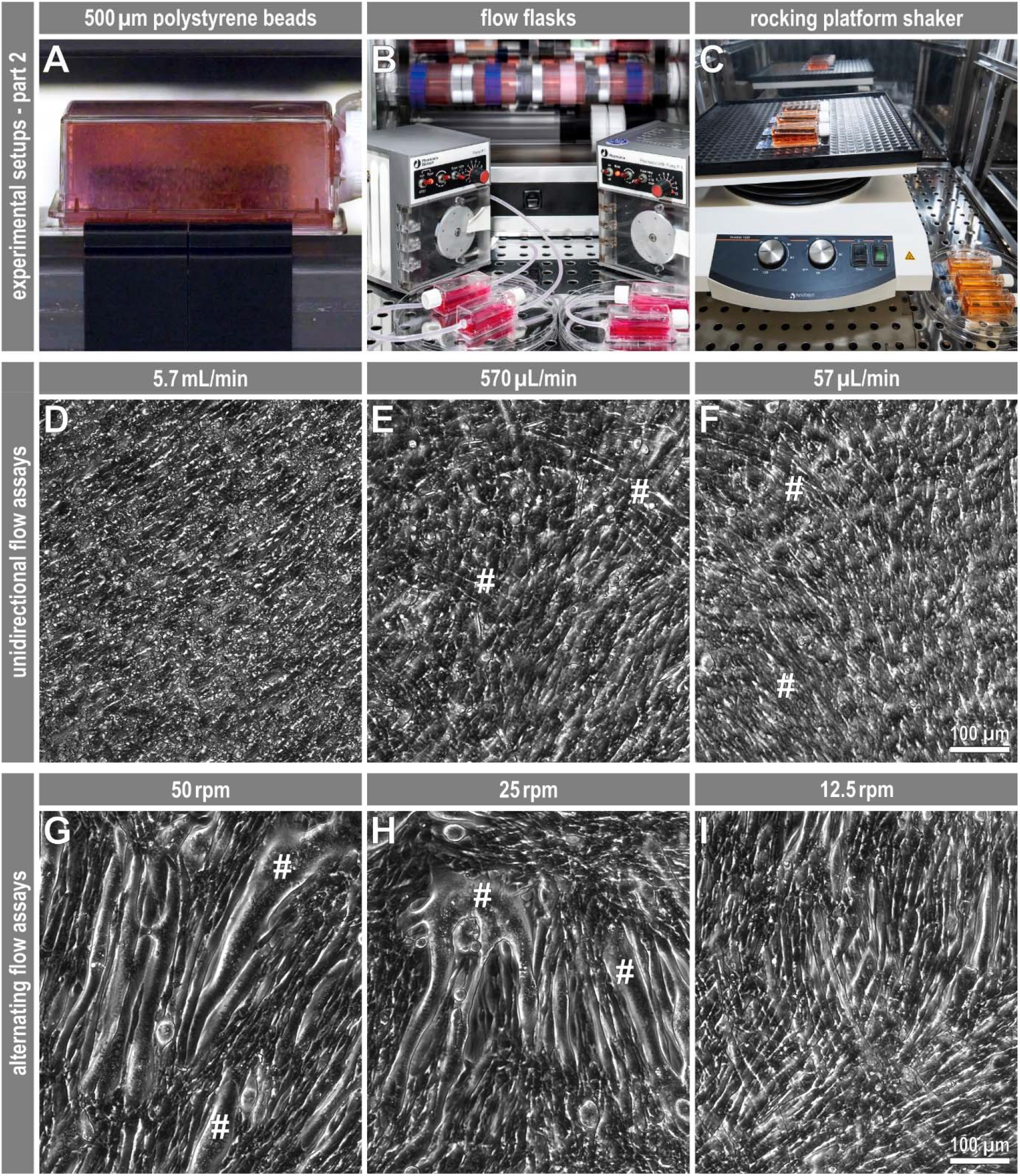
Fluid motion markedly influences C2C12 myotube formation. (**A**) Tests with the addition of 500 μm diameter polystyrene tracer particles to slide flasks filled with medium demonstrate the possible presence of a cylindrical fluid motion during fast 2D-clinorotation. (**B**) Enlarged view of the two peristaltic pumps with the flow flasks connected. (**C**) Slide flasks containing confluent layers of C2C12 myoblasts and differentiation medium are placed on a 5°-tilt-angle rocking platform; non-rocking controls are located to the right of the device. (**D**-**F**) Phase contrast microscopic views of C2C12 cells from the central area of flow flasks perfused with differentiation medium at pump flow rates of 5.7 mL/min (equivalent to an average medium flow speed of 0.24 mm/s within the slide flask; D), 570 μL/min (0.024 mm/s; E), and 57 μL/min (0.0024 mm/s; F) after 6 days of differentiation. The unidirectional medium flow markedly inhibits myotube formation. Hashtags, areas with formation of a few myotubes at the lower flow rates. (**G**-**I**) Phase contrast microscopic views of C2C12 cells from the central area of slide flasks filled halfway with differentiation medium and 5°-tilt-angle rocked at 50 rpm (G), 25 rpm (H), and 12.5 rpm (I). Six days of differentiation under fast to moderate, alternating fluid motion result in the formation of unusually short and wide myotubes; hashtags, large and multinucleated syncytia.

**Figure 3.**
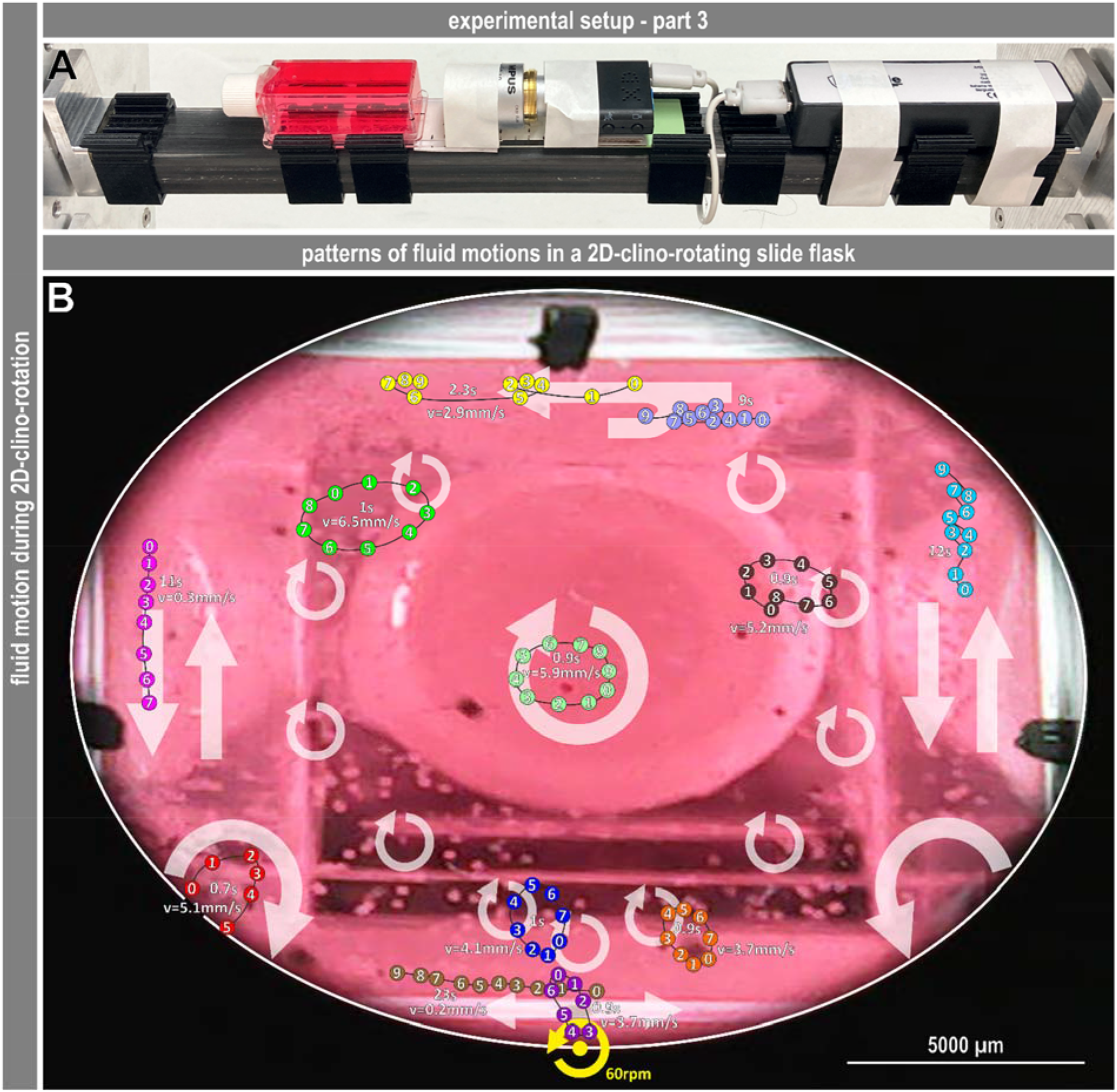
Fluid motion within 2D-clinorotating slide flasks. (**A**) View of the experimental setup for documenting fluid motion in a slide flask during fast 2D-clinorotation. In line from left to right, slide flask filled with differentiation medium and polystyrene tracer particles, a 4x plan objective, a mini-camera (640x480p, 10fps), and a power bank. (**B**) Compilation of the observed flow directions and speeds during the steady-state phase of fast 2D-clinorotation. Background image, exemplary camera view (video snapshot) in longitudinal axis of the slide flask; multiple tracer particles are visible. Grey straight and circular arrows, position and direction of mainly occurring flows (up, down, horizontal, circular) observed with 250 μm or 500 μm whitish-transparent and a few blackened tracer particles. Coloured and numbered dots, tracks of representative individual particles with indication of their mean speeds. Yellow marker, position of the rotational axis at the slide and rotation direction. A total of 83 video sequences were evaluated; an exemplary video clip is available in the supplement (Video S1). Note that since the fluid shear rate (the gradient of the fluid flow speed) is low in the bulk of the medium, the fluid motion is not measurably perturbed by the presence of the beads.

These experiments demonstrated the presence of considerable fluid motion. Observation of the entire particle population as well as tracking of individual particles in different zones of the rotating slide flask during the steady-state phase of fast 2D-clinorotation revealed a large cylindrical fluid motion around the long axis of the side flask, superposed on a complex pattern of flow directions and speeds (Fig. 3B). Close to the cells (100 to 250 μm distance), we observed speeds of particles in a range of 2.7 to 6.5 mm/s, corresponding to shear stresses on the adherent, differentiating myoblasts of around 15 to 25 mPa, assuming a viscosity of the culture medium of 1 mPa s and a linear decrease of the fluid flow speed from the particle to the cell surface (no-slip condition). It has been shown that shear stresses in such a range cause significant cellular responses in e.g., human primary cerebral cortex astrocytes at >50 mPa (Khodadadei et al., 2021), murine primary cortical astrocytes at 25 mPa (Wakida et al., 2020), or C8-D1A murine cerebellar astrocytes (ATCC CRL-2541) at 3 mPa (Lee et al., 2020). Remarkably, human primary osteoblastic cells responded with increased proliferation already to shear stresses as low as 30 μPa (Liegibel et al., 2004). Therefore, the observed suppression of C2C12 myotube formation in a narrow region along the rotational axis may be attributed to shear stress.

To test how myotubes that are not exposed to clinorotation respond to fluid motion, we performed two simple experiments. First, we modified the slide flasks by adding a fluid inlet and outlet along the end faces (Fig. 2B). We then perfused the ‘flow flasks’ with equilibrated (5% CO2, 37°C) differentiation medium using a peristaltic pump at mean flow rates of 57 μL/min, 570 μL/min, and 5.7 mL/min. A flow rate of 5.7 mL/min is equivalent to an average medium flow speed of 0.24 mm/s within the slide flask, and the resulting fluid shear stress at the cell layer is 95 μPa. This flow completely suppressed myotube formation and also caused accumulation of cell debris (Fig. 2D). Similarly, pump flow rates of 570 μL/min (0.024 mm/s medium flow in the flask and 9.5 μPa shear stress at the cells) and even 57 μL/min (0.0024 mm/s, 0.9 μPa) markedly inhibited myotube formation (Fig. 2E, F).

In a second experiment, we placed the slide flasks filled with 10 ml of medium (1 cm fill height) on a rocking platform shaker with a 5° tilt angle to generate an alternating fluid motion along the long axis of the flasks (Fig. 2C). Fast (50 rpm, corresponding to a fluid shear stress of 4.3 mPa (Zhou et al., 2010), and intermediate (25 rpm, 2.1 mPa) fluid motion resulted in the formation of unusually short myotubes with increased width and also larger multinucleate syncytia (Fig. 2G, H), whereas slow (12.5 rpm, 1.1 mPa) fluid motion had no apparent effect (Fig. 2I). Thus, in line with previous reports (Juffer et al., 2014; Jung, 2022), the specific type of fluid motion strongly influences the differentiation of myoblasts into myotubes: While unidirectional, steady, pulsatile and also interval flows suppress myotube formation even at very low rates (Juffer et al., 2014; Jung, 2022), a periodic flow pattern, as shown here, promotes myotube formation. Although these experiments can only approximate some aspects of the complex flow conditions in a rotating clinostat, they show how sensitive myoblasts can respond to different fluid flow patterns.

## Discussion

In this study, we monitored the differentiation process of C2C12 murine myoblasts into multinucleated myotubes under fast 2D-clinorotation. We found impaired myotube formation along a narrow band of approximately 4 mm width near the rotational axis, which has previously been interpreted as being caused by simulated microgravity conditions that are not present off-axis outside this band (Acharya et al., 2022; Brungs et al., 2016; Eiermann et al., 2013; Ivanova et al., 2011). However, offsetting the axis of rotation by 4 mm (‘raising’ the slide surface), which imposes a centrifugal force of 0.016 RCF, still impaired myotube formation along a narrow band near the centreline of the slide. We reasoned that the observed effects may not have been caused by the assumed microgravity but by rotation-induced fluid motion. Indeed, we found that myotube differentiation was impaired by continuous fluid flow. In addition, we observed pronounced fluid motion in the rotating slide flasks, strongly suggesting that fast-rotating 2D-clinostats induce cellular responses that are unrelated to microgravity.

### Cell biological results from fast clinorotation cannot be attributed to microgravity unless alternative explanations have been rigorously tested and excluded

For the vast majority of cell biology experiments, it will be prohibitively expensive to verify findings from ground-based simulated microgravity conditions in true microgravity in space/earth orbit. Therefore, a particularly thorough and critical discussion of ground-based results is needed. Several authors have already cautioned that some findings obtained under simulated microgravity may in fact be attributed to confounding effects such as mechanical vibrations or fluid motion (Herranz et al., 2013; Wüest et al., 2015; Wüest et al., 2017; Zhang et al., 2022). To address these concerns, the mechanical stress-sensitive dinoflagellate *Pyrocystis noctiluca* (NCBI:txid66792), which produces the nocturnal bluish glow of breaking sea waves, was used in a previous study as a reporter for fluid motion and shear stress in a study primarily designed to compare the overall performance of different devices (Hauslage et al., 2017). In that study, a random positioning device was used in 2D-clinostat mode (60 rpm, inner frame), in random speed mode (variable, max. 10 rpm, both frames), in full random mode (random speed, variable, max. 10 rpm, and random direction, both frames), but not in 3D-clinostat mode (e.g., 60 rpm, both frames). *P. noctiluca* in suspension culture responded with less bioluminescence to 60 rpm 2D-clinorotation than to random positioning. This result has been interpreted as a demonstration that a 2D-clinostat can simulate microgravity with negligible fluid motion or shear stresses for suspended dinoflagellates (Hauslage et al., 2017). However, this interpretation was later indiscriminately extended also to cultured mammalian cells, either in suspension or adherent as in this work, without ever testing whether fluid flow and shear stresses are negligible in these settings as well.

Our studies using tracer particles in fast 2D-clinorotating slide flasks demonstrate substantial fluid motion and show that cellular responses to shear stresses cannot be ruled out. Our data experimentally confirm that suppressed C2C12 myotube formation is only present near a narrow band along the rotational axis during clinorotation. This finding has previously been interpreted as i) that microgravity conditions suppress myotube formation, and ii) that perfect microgravity conditions are only present within a 2-3 mm radius around the rotational axis. The latter interpretation is generally accepted and strictly adhered to for sampling. For example, 1F6 human melanoma cells (Fontijn et al., 2009; Van Muijen et al., 1991) grown in slide flasks and 2D-clinorotated at 60 rpm for 24h showed significantly reduced guanylyl cyclase A mRNA levels limited to samples taken from the inner 6 mm wide axial area (Eiermann et al., 2013). Moreover, ML-1 (Schönberger et al., 2000) and RO82-W-1 (Estour et al., 1989) follicular thyroid cancer cells grown in slide flasks and 2D-clinorotated at 60 rpm for either 3 or 7 days showed marked formation of actin stress fibres and subplasmalemmal enrichment of F-actin within the inner 6 mm wide axial area (Svejgaard et al., 2015). Another comprehensive study in spontaneously beating cardiomyocytes derived from human-induced pluripotent stem cells grown in modified slide flasks and 2D-clinorotated at 60 rpm for 2 days showed multiple mitochondria-, contraction-, and senescence-related changes and dysfunctions in samples taken from the inner 3 mm of the axial region. Notably, that study also reported the formation of actin stress fibres and sarcolemmal caveolae in the fast 2D-clinorotated cardiomyocytes as a clear indication of the presence of mechanical stress (Acharya et al., 2022).

However, the common notion that cellular effects due to simulated microgravity are confined only to a narrow region close to the rotational axis is contradicted by our findings. In particular, when we eccentrically rotated our slides 4 mm off-axis, we still observed the same narrow band of reduced myotube formation. Impaired myotube formation was also reported for 60 rpm 3D-clinorotation of differentiating C2C12 myoblasts (Calzia et al., 2020). For such 2-axis rotation, substantial fluid motion and shear stress were reported even at lower rotational speeds (see above), so that a lower rotational speed was recommended to reduce fluid motion and shear stress (Wüest et al., 2017; Zhang et al., 2022). Considering that fluid motion is a potent stimulus for numerous cell responses – which we also demonstrate in our present study – previously published results obtained with fast-rotating clinostats in combination with adherent mammalian cells, a selection of which are cited here, need to be reinterpreted. While we do not question the validity of the reported data, we caution that the reported cellular effects observed in fast-rotating clinostats are most likely not due to simulated microgravity but rather to fluid flow effects. The need for a critical interpretation of these results is further underscored by the fact that even a fluid flow or shear stress roughly a thousand times smaller (≈30 μPa) than those we observed (15-25 mPa) can cause substantial cellular effects (Liegibel et al., 2004).

A recent study using pre-formed myotubes from L6 rat myoblasts (ATCC CRL-1458) compared changes that occurred due to i) real microgravity versus 1 xg control condition produced by centrifugation in the space station, and ii) ground-based, random-mode 3D-clinorotation versus static, non-rotating 1 xg control condition (Uchida et al., 2018). Although this work is not directly comparable to ours, the study reported reduced myotube diameters after 10 days in the space station and after 3 days in random-mode 3D-clinorotation. However, the ground-based static non-rotating, ‘true 1 xg control’ and the ‘1 xg centrifugation control’ in the space station resulted in markedly different myotube diameters (approximately 5 μm between both controls, see figure 2A, B in (Uchida et al., 2018)). This difference is three to four times higher than the differences between the two experiment/control pairs. Thus, we argue that the cell responses under real microgravity versus ground-based simulated microgravity are not directly comparable because they are likely to have different underlying mechanisms.

### Strategies for improved experimental design

To identify spurious cellular responses caused by effects other than the desired simulated microgravity, it is necessary to include further control experiments in addition to static, non-rotating cell samples, for example offset-rotating controls as used in the present work, or a vertical-rotating control (see Supplementary Fig. 1 in (Li et al., 2022), and (Gruener and Hoeger, 1991)). A vertically rotating control introduces a gravitational vector that is no longer perpendicular to the cell-matrix interface, and therefore requires a corresponding vertical static, non-rotating culture as a further control. Additionally, other controls should be included to specifically assess potential effects of fluid motion, depending on the individual sensitivity of the cell type used, which can vary considerably.

Further, it is paramount to reduce or altogether prevent the intrinsic, rotation-induced fluid flow. To accomplish this, it is first necessary to understand the cause of the fluid moving in a 2D-clinostat, which is not obvious. In fact, previous fluid dynamic modelling suggested that the culture medium in a 2D-clinorotating culture flask becomes steady after an initial phase of motion (Klaus et al., 1998; Wüest et al., 2017). However, this is only true for ideal conditions, in particular for a perfectly steady rotational speed, and for a perfectly steady axis of rotation. In practice, small fluctuations of the drive motor, combined with small oscillatory lateral movements due to the eccentric rotation of the culture vessels, will inevitably cause fluid motion due to inertial and Coriolis forces.

A first, simple strategy is therefore to reduce the circulating fluid mass by reducing the height of the slide flask or culture vessel, but this would require a gas-permeable flask material and would also limit the duration of the experiment before the cell culture medium needs to be exchanged. One option would be to use smaller culture vessels, such as channel slides, where the medium is cyclically exchanged at regular time intervals by connected pumps and reservoirs. However, vessels of very small dimensions can also influence the cells, so that the influence of the geometry of the vessels and their surface properties, including any surface micropatterning, would also have to be taken into account. Another strategy could be to increase the viscosity of the cell culture medium. This would reduce the speed of fluid motion, but at the same time the positive effect could be cancelled as the fluid shear stress at the cell surface increases in proportion to the viscosity. The latter problem could be avoided by completely filling the culture vessel with an elastic hydrogel as culture medium. This hydrogel would need to be sufficiently permeable for gas exchange and nutrient transport. Possible examples are inert biopolymer-based hydrogels on the basis of alginate or agarose. Furthermore, matrix protein-based hydrogels containing e.g., collagen, fibrin, or Matrigel could offer the possibility to explore cell or microtissue behaviour under true 3D-culture conditions in combination with simulated microgravity. At the same time, the technical design of a clinostat must be improved to minimize rotational speed fluctuations and to maximize the stability of the rotational axes.

Because the rotating slide flasks in the 2D-clinostat are filled with 2 cm of medium, the hydrostatic pressure acting on the cells changes sinusoidally at a frequency of 1 Hz between 0 and 2 cm H2O (200 Pa), minus a small pressure originating from the outward-acting centrifugal force. While it seems quite unlikely that the cells would respond to pressure changes of such a small magnitude (Kao et al., 2017; Purkayastha et al., 2021; Yu et al., 2013), a hydrostatic pressure of 200 Pa may cause small deformations of the plastic substrate on which the cells were grown, and therefore may induce measurable cell responses. The application of a constant versus a sinusoidally varying hydrostatic pressure should therefore be considered as a further control experiment.

But even with a perfect clinostat design that completely prevents fluid flow and shear stresses, experimenters should question the assumption that microgravity-like conditions can be achieved by averaging-out the vector of gravity. For example, bone cells strongly respond to oscillatory fluid shear stress even when the stress has a time-average of zero (Ponik et al., 2007). On an even more fundamental level, researchers should also question the assumption that gravitational forces are of relevant magnitude to be sensed by a single cell in a near-buoyant, aqueous environment. Considering that C2C12 cells as used in this study generate traction stresses of typically 1 kPa or more (Sakar et al., 2012), normal gravitational stresses acting at the cell-matrix interface are more than 5 orders of magnitude smaller. In fact, the total gravitational forces of a single cell in water of around 1 to 2 pN (assuming a cell density of 1.05 g/cm^3^) is smaller than the contractile force generated by a single myosin molecule. Thus, apart from the unsolved technical challenges, the problem of separating the cellular effects of these extremely small gravitational forces from the combined effects of much larger contractile, thermal, and fluid forces or geometric constrains remains to be addressed. This could be achieved by comprehensive transcriptomic or proteomic analyses of samples from different experimental conditions e.g., true microgravity, simulated microgravity by clinorotation, fluid flow, and vibration, combined with multiple-comparison data analysis of the resulting omics-patterns. Consideration of the strategies discussed here for improved assay design requires further studies and inevitably adds complexity to the interpretation of results.

### Clinorotation and ‘simulated microgravity’

The term ‘simulated microgravity’, now commonly used to describe ground-based laboratory conditions as a workaround for experiments in real microgravity, should not be taken to mean that gravitational forces have disappeared or been eliminated. In this respect, 2D-clinorotation should be understood as an attempt to mimic certain aspects of microgravity, but not as a strategy to simulate it in its entirety. Therefore, for each cell type, clinostat device, and experimental setting, it remains important to clearly define and then verify which aspect of microgravity can be realistically mimicked. This can be a time-consuming process. Final verification of a result obtained in this way can only be done in real microgravity. However, this does not eliminate the need for close observation and extensive controls to identify deviations, side effects and confounding variables in a ground-based clinostat experiment. And even in real microgravity, efforts must be made to distinguish between putative cellular effects of gravitational forces and, for example, thermal molecular forces, electrostatic forces between the cell surface and the substrate, or the contractile force of the cellular actomyosin machinery.

## Methods

### Exposure of differentiating C2C12 muscle cells to fast 2D-clinorotation and fluid motion

C2C12 myoblasts derived from C3H mouse skeletal muscle (ECACC 91031101, also available as ATCC CRL-1772) with low passage number (highly myogenic passage no. 6 at start of this work) were grown in DMEM supplemented with 10% fetal calf serum, 2 mM L-glutamine, 1 mM pyruvate, 1x non-essential amino acids, 100 U/ml penicillin, and 0.1 mg/ml streptomycin (all from Pan Biotech, Aidenbach, Germany) in a NuAir incubator (NU-5841E; ibs tecnomara GmbH, Fernwald, Germany) at 5% CO2 and 37°C. Semi-confluent cultures were split using 0.05% trypsin/0.5 mM EDTA (Pan Biotech, Aidenbach, Germany) by 1:8 for assays in SlideFlasks (#170920; Nunc/Thermo Scientific, Waltham, USA) or 1:20 for maintenance in standard 100 mm culture dishes (#83.3902; Sarstedt, Nümbrecht, Germany). Differentiation of the myoblasts into myotubes was induced by replacing the growth medium with a low-mitotic medium i.e., the former growth medium with the fetal calf serum replaced by 5% horse serum (Gibco/Thermo Scientific, Waltham, USA). The used horse serum was from a batch that had been tested regarding muscle cell differentiation. Myotube formation was monitored up to 8 days.

For exposure to fast 2D-clinorotation, C2C12 myoblasts were grown to confluency in slide flasks within 1 day after splitting, and immediately after addition of differentiation medium clinorotation was done for 6 days. The culture vessels were attached to a custom-made slide flask-2D-clinostat (Eiermann et al., 2013; Mansour Jamaleddine, 2021) shelved in the above-mentioned incubator at 5% CO2 and 37°C (Fig. 1A-C); static, non-rotating control slide flasks were placed on top of the clinostat frame to rule out the possibility of effects of the unavoidable device vibration (Fig. 1A). Rotation speed was 60 rpm (360 deg/s, 6.3 rad/s) generating few millimetres wide central band-formed areas of centrifugal acceleration, for example (band width, acceleration, fraction of Earth’s gravity): 3 mm, 0.059 m/s^2^, 0.006 RCF; 4 mm, 0.079 m/s^2^, 0.008 RCF; 6 mm, 0.118 m/s^2^, 0.012 RCF; 8 mm, 0.158 m/s^2^, 0.016 RCF; 12 mm, 0.237 m/s^2^, 0.024 RCF; 16 mm, 0.316 m/s^2^, 0.032 RCF; 18 mm, 0.355 m/s^2^, 0.036 RCF. In addition, sets of culture vessels were attached in a way that the slide was positioned 4 mm above the rotational axis; this was achieved by holding clamps with a thicker intermediate base (Fig. 1B, blue clip vs. black clip, and axis of rotation label in yellow). Clinorotation experiments were independently performed three times with four slide flasks for each condition (clino-rotation, offset-rotation, static non-rotating control). All images and data are from cells located within a central 4 mm wide area along the rotational axis across the entire slide. Clinorotating slide flasks were completely filled with incubator-equilibrated medium, carefully sealed with Parafilm M (Bemis, Neenah, USA) and the screw caps, and checked for total absence of air bubbles during the entire duration of the experiment. All ‘Bonn criteria’ for a clinostat device (Hammond and Allen, 2011) are covered by parameters specified in the two above paragraphs.

For 5 to 7 days exposure to unidirectional flow, C2C12 myoblasts in custom-modified slide flasks, termed ‘flow flasks’ (Jung, 2022), were grown to confluency within 1 day after splitting. Immediately after addition of differentiation medium, culture vessels were connected to peristaltic pumps (Pump-P1; Pharmacia Biotech AB, Uppsala, Sweden) using 2 mm inner diameter PBS-washed and autoclaved silicone tubes (Fig. 2B). Pump flow rate was set to either 10x10 (5.7 mL/min), 1x10 (570 μL/min), or 1x1 (57 μL/min); exact flow rates were determined by measuring 1-minute-flow volumes at the end of each experiment. Flow experiments were independently performed three times (one flow setting per experiment) with four slide flasks for each condition (flow and static control). Images were obtained from cells located within a central 4 mm wide area across the entire long axis of the slide.

For 5 to 7 days exposure to oscillating flow along the long axis of slide flasks, C2C12 myoblasts were also grown to confluency within 1 day after splitting. Immediately after the addition of differentiation medium, the slide flasks were centrally placed on a 5°-tilt-angle rocking platform shaker (Duomax 1030; Heidolph, Kelheim, Germany) (Fig. 2C). Rocking rate was set to either fast (50 rpm), medium (25 rpm), or slow (12.5 rpm) and verified by counting the 1-minute-number of cycles. Rocking experiments were independently performed three times (one rocking rate per experiment) with three slide flasks for each condition (rocking and static control). Images were obtained from cells located within a central 4 mm wide area across the entire long axis of the slide.

### Microscopic imaging, fluid motion monitoring, and data analysis

Phase contrast microscopic images were recorded with a transmission light microscope (DMIL LED; Leica Microsystems, Wetzlar, Germany), objective HI PLAN I 10x/0.22 PH1, equipped with a heated stage (Tempcontrol-37; Pecon, Erbach, Germany) and stand-alone FLEXACAM C1 colour camera (Leica Microsystems, Wetzlar, Germany). Using the phase contrast images in random order, myotubes were independently assessed and counted by two expert authors of this study. The widths of these myotubes was measured at their position of maximal width and respective diameters were determined using ImageJ v.1.53 (Schneider et al., 2012). Statistical analysis was performed using Prism v.6.07 (GraphPad Software Inc., San Diego, USA); data passed D’Agostino & Pearson omnibus normality testing, and statistical significances were calculated using one-way ANOVA with Tukey’s multiple comparisons correction and subsequent unpaired two-tailed Student’s t-tests. Data are shown as scatter dot plots with medians and interquartile ranges. For fluid motion monitoring, a mini-camera (no brand, 640x480 pixel resolution at 10 fps) complemented with an objective (Olympus, 4x/0.10, CX22 PL4X) was fixed facing the front side of a slide flask (Fig. 3A). Video frame snapshots were created using VLC media player v.3.0.18 (VideoLAN Organization) and used to manually generate single-bead-tracks. The supplementary video clip was generated using ClickPoints (Gerum et al., 2016). Images were processed and figures assembled using CorelDraw Graphics Suite X7 (Corel Corporation, Ottawa, Canada).

## Declarations

### Authors’ contributions

J.M., C.B., and C.S.C. jointly conceived the study, reviewed data, designed figures, and drafted the manuscript. J.M., C.B., M.J., and C.S.C. designed and carried out experiments and analysed data. J.M. and C.S.C. performed statistical evaluations. L.E., B.F., and C.S.C. analysed data, reviewed data, and jointly finalised the manuscript; C.S.C. prepared the final version of manuscript figures. All authors have read and agreed to the final version of the manuscript.

## Acknowledgements

We gratefully acknowledge the following support of this work: Dr. Philip Born, Institute of Materials Physics in Space, German Aerospace Center, Cologne, Germany, for providing polystyrene beads; Mr. Sebastian Steinhäuser and Dr. Thomas Voigtmann, Institute of Materials Physics in Space, German Aerospace Center, Cologne, Germany, for helpful discussions of fluid motion profiles; Dr. Peter van der Ven, Institute for Cell Biology, University of Bonn, Germany, for providing horse serum from a batch that had been tested for C2C12 muscle cell differentiation; and last but not least Mr. Marvin Diegeler, Photo Services Cologne, German Aerospace Center, Cologne, Germany, for professional photography of experimental setups. We would also like to thank the three Referees of this work, whose expertise, attention to detail and extraordinary dedication have contributed to the maturity of this work.

## Data availability

The data that support the findings of this study are available within the article and its supplementary material.

## Supplementary materials

**Supplementary video 1. Fluid motion within 2D-clinorotating slide flasks.** Exemplary video clip documenting marked fluid motion in a slide flask during the steady-state phase of fast 2D-clinorotation. Multiple such video clips were used to compile the complex pattern of observed flow directions and speeds as shown in figure 3.

